# B cells from aged mice do not have intrinsic defects in affinity maturation

**DOI:** 10.1101/2023.04.24.538044

**Authors:** Jia Le Lee, Silvia Innocentin, Alyssa Silva-Cayetano, Stephane M. Guillaume, Michelle A. Linterman

## Abstract

Affinity maturation, the progressive increase in serum antibody affinity after vaccination, is an essential process that contributes to an effective humoral response against vaccines and infections. Germinal centres (GCs) are key for affinity maturation, as they are where B cells undergo somatic hypermutation of their immunoglobulin genes in the dark zone, before going through positive selection in the light zone via interactions with T follicular helper cells and follicular dendritic cells. In aged mice, affinity maturation has been shown to be impaired, but whether B cell-intrinsic factors contribute to this defect remains unclear. In this study, we show that B cells from aged B cell receptor transgenic mice are able to become GC B cells, which are capable of receiving positive selection signals to a similar extent as B cells from young adult mice. Consistent with this, ageing also does not impact the ability of B cells to undergo somatic hypermutation and acquire affinity-enhancing mutations. Together, this shows that there are no B cell-intrinsic defects in affinity maturation with age when the B cell receptor repertoire is constant.

## Introduction

Affinity maturation, the progressive increase in affinity of serum antibodies over time, is an important process that underlies an effective humoral response against vaccines and infections (Eisen & Siskind, 1964; Nussenzweig & Benacerraf, 1967). Germinal centres (GCs) are the cellular engines of affinity maturation. Within GCs, B cells undergo somatic hypermutation of their immunoglobulin genes in the dark zone, which generates a pool of B cells carrying random mutations that will then undergo selection in the light zone (Mesin et al., 2016). B cells carrying functional B cell receptors (BCRs) that can uptake and present antigen to T follicular helper (Tfh) cells will receive positive selection signals, which induces upregulation of the proto-oncogene cMyc, and promotes cyclic reentry in the dark zone for further somatic mutations and clonal expansion (Calado et al., 2012; Dominguez-Sola, Victora, Ying, Phan, Saito, Nussenzweig, et al., 2012; Victora et al., 2010). Eventually, these B cells exit the GC as memory B cells or long-lived antibody-secreting plasma cells, which are key in conferring protection against future infections.

The impaired vaccine response during ageing has been widely characterized across different vaccine formulations. This defect involves not only a quantitative reduction in vaccine-specific antibody titres (Collier et al., 2021; Hill et al., 2022; Li et al., 2017), but also higher incidence of non-specific autoantibodies (Howard et al., 2006) and reduced accumulation of productive *de novo* immunoglobulin mutations to adapt to drifted virus strains in older individuals (Henry et al., 2019). Analysis of the immunoglobulin genes of GC B cells from the Peyer’s patches and spleens of older people revealed that the mechanism of somatic hypermutation is unaltered with age, but that the selection mechanisms may be altered (Banerjee et al., 2002). *In vivo* studies in mice show that affinity maturation is impaired with age, as shown by fewer high-affinity GC B cells and fewer antigen-specific plasma cells in the bone marrow of aged mice post-vaccination (Han et al., 2003; Silva-Cayetano et al., 2023; Yang et al., 1996). This age-related defect in the GC response and its output is a result of a reduction in GC magnitude as well as impairment in the selection process (Han et al., 2003; Miller & Kelsoe, 1995; Silva-Cayetano et al., 2023; Yang et al., 1996).

Since an effective humoral response during vaccination relies on the coordinated interaction of multiple cell types, a multitude of factors can contribute to age-related defects in vaccine responses. Previous studies have shown that B cells from older people and aged mice have no intrinsic defects in responding to stimulation and differentiating into plasma cells (Dailey et al., 2001; Lee et al., 2022; Yang et al., 1996). However, whether there are B cell-intrinsic defects in the process of affinity maturation remains unclear. In this study, we tracked the response of antigen-specific B cells derived from 6-12 weeks old younger adult and >90 weeks old aged B1-8i B cell receptor transgenic mice, transferred into young recipient mice, over time following immunisation. B cells from aged mice had no defects in undergoing class-switch recombination, nor in becoming GC B cells or plasmablasts, compared to those from young adult mice. We show that B cells derived from aged B1-8i mice were equally able to upregulate cMyc in the GC, suggesting no intrinsic defects in their ability to receive positive selection signals. Finally, sequencing of the V_H_186.2 heavy chain region of NP-specific GC B cells derived from young and aged B1-8i mice revealed no age-related intrinsic defects in the rate of somatic hypermutation nor their ability to acquire the affinity-enhancing W33L mutation. This highlights that the poor quality of the GC response in ageing is not due to B cell-intrinsic impairments.

## Results

### B cells in aged mice are less frequently of a follicular phenotype

The B1-8i adoptive transfer system was used in this study to interrogate if there are any cell-intrinsic defects in the ability of B cells from aged mice to undergo affinity maturation. About 10% of B cells from B1-8i mice contain the knock-in canonical B1-8 heavy chain (V_H_186.2, DFL16.1 and J_H_2) that, when combined with an Igλ light chain, produces an antibody with intermediate affinity for the hapten 4-hydroxy-3-nitrophenylacetyl (NP) (Shih et al., 2002). In order to check for phenotypic differences that might affect B cell responses to immunisation, NP-specific B cells from young adult (6-12 weeks old) and aged (>90 weeks old) B1-8i mice were first stained with a comprehensive flow cytometry panel to check for their Ig isotype and basal expression levels of markers prior to transfer. There was no significant difference in the percentage of IgD+ naïve B cells among NP-specific B cells from young adult versus aged donor mice **(Fig. 1a-b)**. However, aged mice tended to have about 10% fewer follicular B cells (CD23hi CD21int) among NP-specific B cells (average of 58%), compared to young adult mice (average of 71%) **(Fig. 1a-b)**. We also assessed the expression of various proteins on the NP-specific B cells: chemokine receptors and trafficking molecules, CXCR4, CD62L and CXCR5 **(Fig. 1c-e)**, B cell activation markers, CD38, GL7 and IRF4 **(Fig. 1f-h)**, atypical or age-associated B cell markers, T-bet and CD11c (Rubtsov et al., 2011) **(Fig. 1i-j)**, costimulatory molecules, CD86, CD40 and MHCII **(Fig. 1k-m)**, and CD11b, which has been shown to be expressed in a subset of regulatory B cells (Liu et al., 2015) **(Fig. 1n)**, and observed no age-dependent differences.

**Figure 1.**
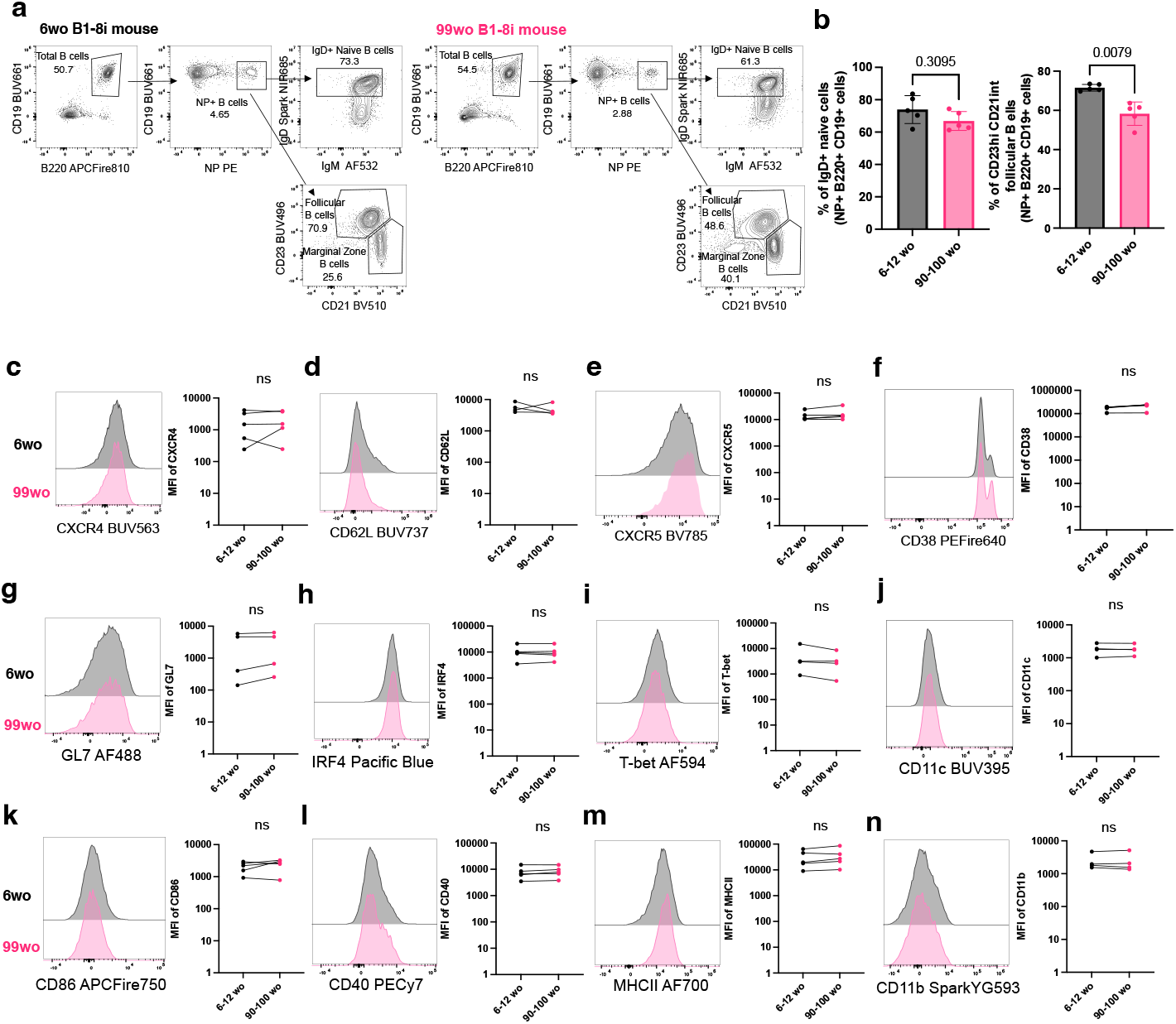
NP-specific B cells from aged mice are phenotypically similar to those from young adult mice. **(a)** Gating strategy for NP-specific IgD+ cells and follicular cells (CD23hi CD21int) in young (6wo) and aged (99wo) B1-8i transgenic mice. Numbers adjacent to gates indicate percentage of parent population. Cells were pre-gated for live cells and single cells. **(b)** Graphs depicting the percentage of IgD+ cells (left) and follicular (CD23hi CD21int) cells (right) among NP+ B220+ CD19+ cells in young (6-12wo) and aged (90-100wo) B1-8i transgenic mice. Bar height corresponds to the mean, error bars indicate standard deviation, and each symbol represents values from individual mice. Statistics were calculated using unpaired Mann-Whitney U test. Data pooled from five independent repeat experiments **(c-n)** Representative flow cytometric histograms and graphs showing the mean fluorescence intensities (MFI) of **(c)** CXCR4, **(d)** CD62L, **(e)** CXCR5, **(f)** CD38, **(g)** GL7, **(h)** IRF4, **(i)** Tbet, **(j)** CD11c, **(k)** CD86, **(l)** CD40 and **(m)** CD11b of NP+ B220+ CD19+ cells from young (6wo) or aged (99wo) B1-8 transgenic mice. Each symbol represents values from individual mice, with young and aged donors from the same experiment shown as paired values. Statistics were calculated using Wilcoxon matched pairs signed rank test. Data pooled from at least four independent repeat experiments.

### NP-specific B cells from aged mice do not have intrinsic defects in becoming GC B cells

To determine if there are any cell-intrinsic defects in B cells from aged mice when responding to stimulation, equal numbers of NP-specific B cells from either a young adult or aged B1-8i mice were adoptively transferred into young (8-12 weeks old) congenic wild-type recipient mice. Their responses were assessed in draining inguinal lymph nodes (iLNs) of recipient mice at days 6, 10, 14, 21 and 28 post-immunisation with NP conjugated to the Keyhole Limpet Hemocyanin protein (NP-KLH) in Alum **(Fig. 2a)**. Across all timepoints, there was no significant difference in the percentage and number of NP-specific B cells derived from the young adult or aged donor mice in recipient iLNs **(Fig. 2b)**. NP-specific B cells from aged mice also had no significant defect in class-switch recombination to IgG1 **(Fig. 2c-d)** or in becoming GC B cells **(Fig. 2e-f)**. There was also no significant difference in the Dark Zone:Light Zone (DZ/LZ) ratio of the donor-derived NP-specific GC B cells from young adult or aged mice **(Fig. 2g-h)**, nor in the percentage and number of short-lived extrafollicular plasma cells derived from donor cells from young adult or aged mice **(Fig. 2i-j)**. Together, this shows that there are no cell-intrinsic defects in the ability of NP-specific B cells from aged B1-8i mice in responding to stimulation and differentiating into plasma cells or GC B cells.

**Figure 2.**
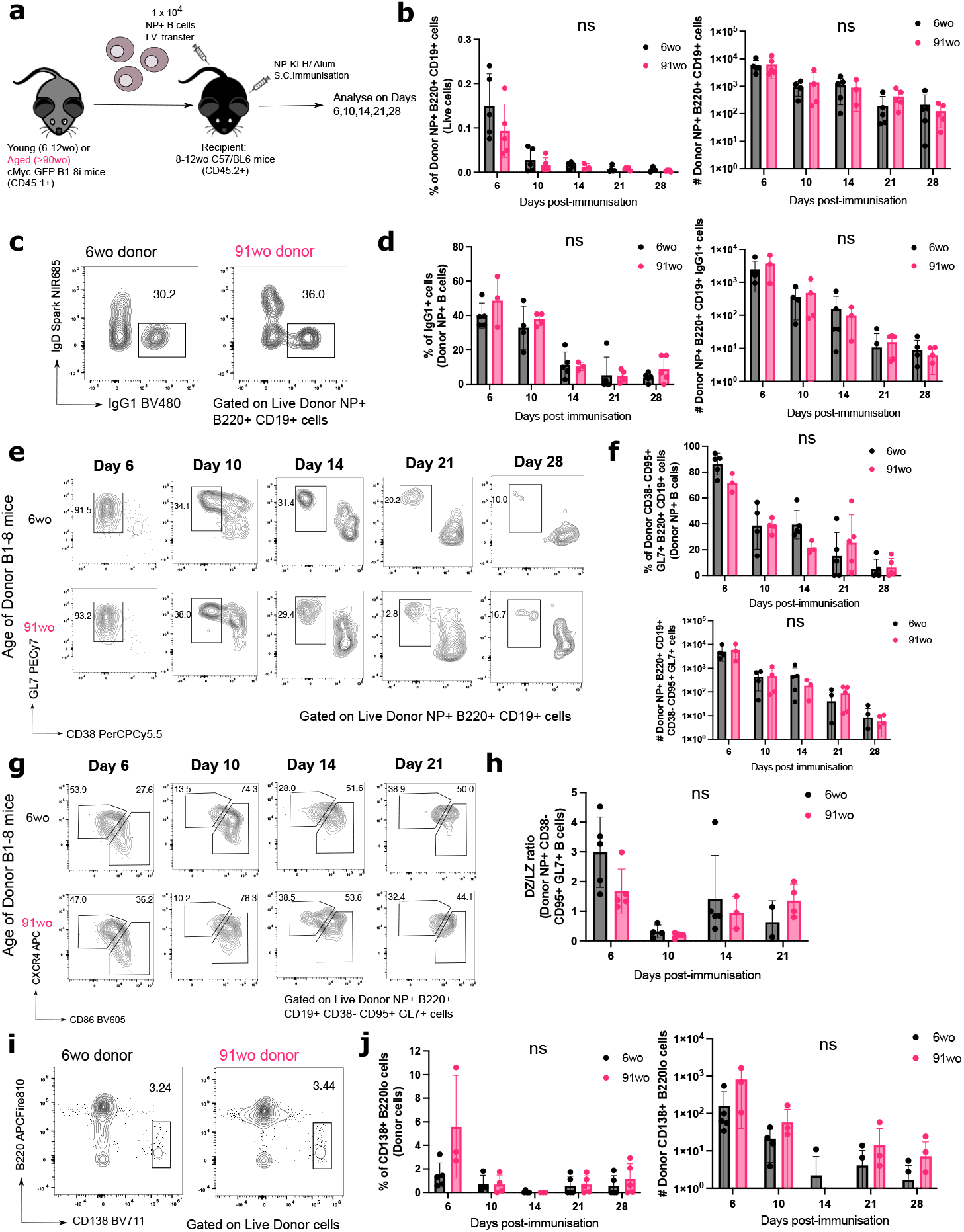
B cells from aged donor mice have no defects in class-switch recombination, entering the GC response and plasmablast differentiation. **(a)** Schematic diagram of adoptive transfer experiments to compare the intrinsic function of B cells from young and aged mice in young recipient mice post-immunisation. 1 × 10^4^ NP-binding CD45.1+ B cells from either a young (6-12wo) or aged (>90wo) B1-8i mouse was transferred intravenously into young (8-12wo) CD45.2+ C57/BL6 SJL mice. The recipient mice then received subcutaneous immunisation with NP-KLH in alum. Draining inguinal lymph nodes (iLNs) were taken at indicated times post-immunisation for flow cytometric and downstream analyses. **(b)** Graphs depicting the percentage and number of donor NP+ B220+ CD19+ cells out of live cells in recipient iLNs at different timepoints post-transfer and immunisation. **(c-f)** Representative flow cytometric plots showing gating strategies for donor-derived IgG1+ IgD-B cells **(c)** and CD38-GL7+ GC B cells **(e)** from 6wo or 91wo donor mice at different timepoints post-transfer and immunisation. Numbers adjacent to gates indicate percentage of donor NP+ B220+ CD19+ cells. Graphs depicting the percentage and number of donor NP+ IgG1+ cells **(d)** and donor NP+ CD38-CD95+ GL7+ cells **(f)** in recipient iLNs at different timepoints post-transfer and immunisation. **(g)** Representative flow cytometric plots of dark zone (CXCR4+CD86lo) and light zone (CXCR4loCD86+) donor-derived GC B cells from 6wo or 91wo donor mice. Numbers adjacent to gates indicate percentage of donor NP+ B220+ CD38-CD95+ GL7+ GC B cells. **(h)** Graphs comparing dark zone: light zone (DZ/LZ) ratio among donor NP+ GC B cells derived from 6wo and 91wo mice. **(i)** Representative flow cytometric plots showing gating strategies for donor-derived CD138+ B220lo cells from 6wo or 91wo donor mice at day 6 post-transfer and immunisation. Numbers adjacent to gates indicate percentage of donor cells. **(j)** Graphs depicting percentage and number of donor CD138+ B220lo cells in recipient iLNs at different timepoints post-transfer and immunisation. Bar height corresponds to the mean, error bars indicate standard deviation, and each symbol represents values from individual recipient mice. Statistics were calculated using 2-way ANOVA with Sidak’s multiple comparisons test. Data representative of two independent repeat experiments.

### NP-specific B cells from aged mice receive positive selection signals in the light zone

Positive selection is a key process that primarily occurs in the LZ of the GC, and it is essential for affinity maturation. Using the aforementioned adoptive transfer system **(Fig. 2a)**, the response of transferred B1-8i B cells, which carry a cMyc-GFP reporter gene, was assessed at the peak of the GC response, day 6 post-immunisation **(Fig. 2e-f)** (gating strategy shown in **Fig. 3a)**. The cMyc+ GC B cells identified were predominantly of the LZ phenotype (CD86hi CXCR4+ GC B cells) (Dominguez-Sola, Victora, Ying, Phan, Saito, Dalla-Favera, et al., 2012) **(Fig. 3b)**. NP-specific GC B cells derived from aged mice showed no intrinsic defects in cMyc upregulation **(Fig. 3c-d)**, suggesting that age does not diminish the ability of B cells to receive positive selection signals in a young microenvironment. This suggests that this mechanism that underpins affinity maturation is intact.

**Figure 3.**
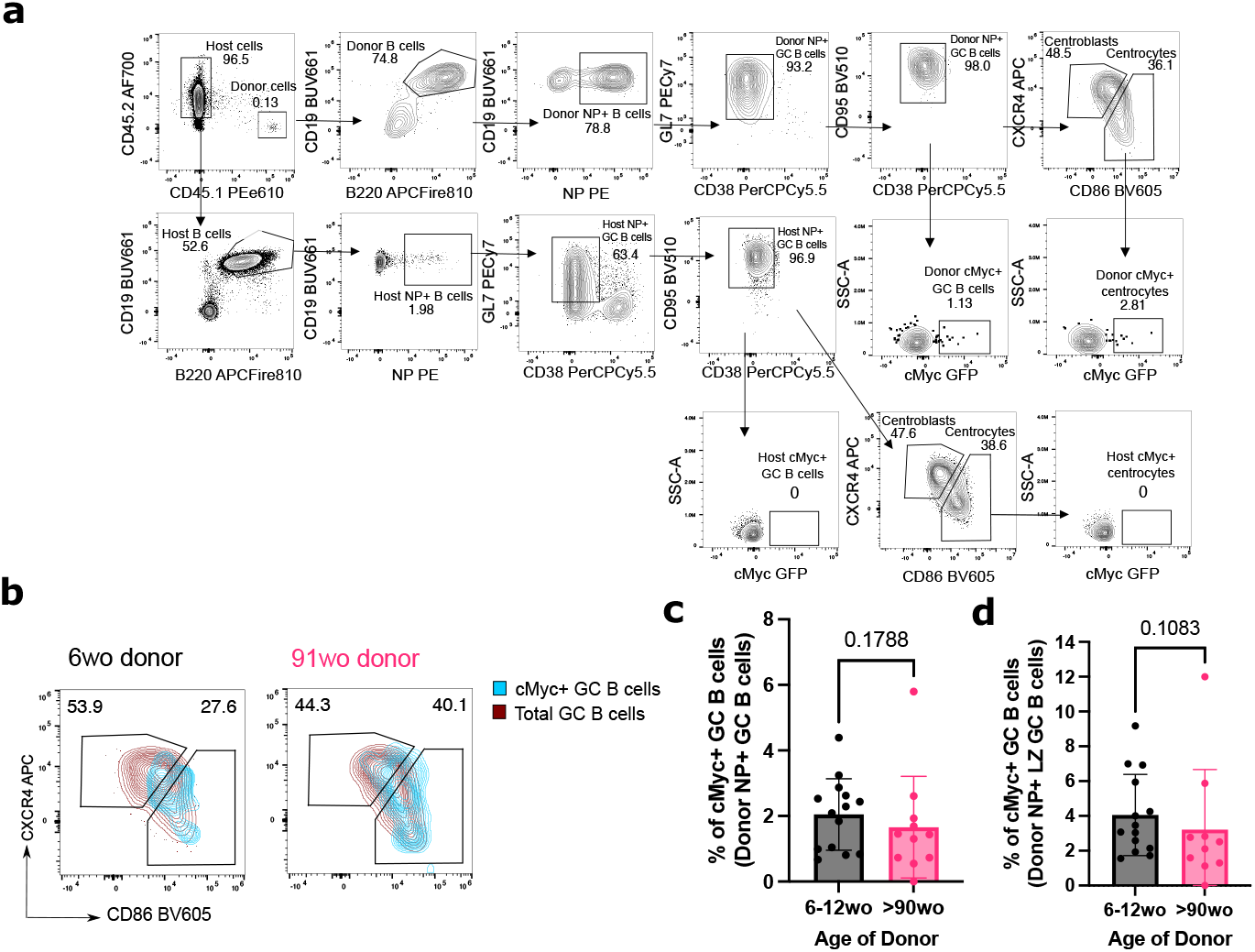
GC B cells from aged donor mice have no intrinsic defects in cMyc expression. **(a)** Representative flow cytometric plots showing the gating strategy for cMyc+ cells from donor GC B cells (CD45.1+ CD19+ B220+ NP+ CD38-GL7+ CD95+) and centrocytes (CXCR4lo CD86hi). **(b)** Flow cytometric plots showing the overlay of cMyc+ GC B cells (blue) on total GC B cells (red) gated for dark zone (CXCR4+CD86lo) and light zone (CXCR4loCD86+) phenotype. Numbers adjacent to gates indicate percentage of donor NP+ CD19+ B220+ CD38-GL7+ CD95+ GC B cells. **(c)** Graphs depicting the percentage of cMyc+ GC B cells out of donor NP+ GC B cells (left) and donor NP+ LZ GC B cells (right) in recipient iLNs at day 6 post-transfer and immunisation. Bar height corresponds to the mean, error bars indicate standard deviation, and each symbol represents values from individual recipient mice. Statistics were calculated using unpaired Mann-Whitney U test. Data pooled from three independent repeat experiments.

### NP-specific B cells from aged mice undergo mutation and selection of high affinity clones

Finally, the ability of B cells from aged mice in undergoing somatic hypermutation and acquiring high-affinity mutations was investigated, by sequencing the V_H_186.2 region of sorted donor-derived NP-specific GC B cells from the iLNs of recipient mice at day 14 post-immunisation. Substitution of a tryptophan (W) with a leucine (L) at position 33 of the complementarity-determining region 1(CDR1) of V_H_186.2 results in a tenfold increase in antibody affinity for NP (Allen et al., 1988). No significant difference was shown in the frequency of W33L mutation among sorted NP-specific IgG1+ GC B cells from young adult versus aged mice **(Fig. 4a-b)**. In addition, the frequency of mutations and the ratio of replacement to silent mutations were similar among GC B cells from young adult and aged mice **(Fig. 4c-d)**. Together, this suggests that B cells from aged mice are equally functional in undergoing somatic hypermutation and receiving positive selection signals, and ultimately producing GC B cell clones with affinity-enhancing mutations.

**Figure 4.**
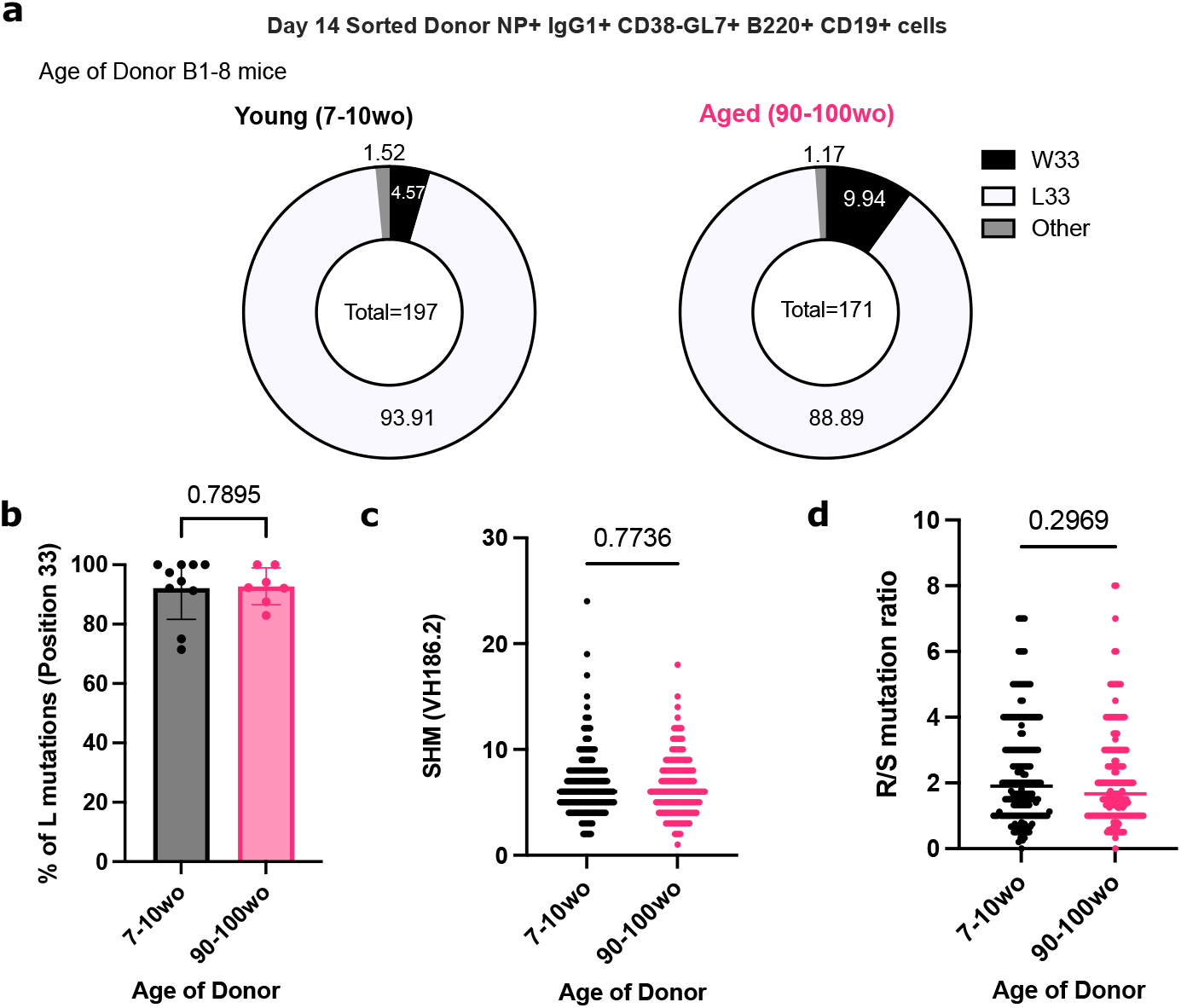
NP-specific B cells from aged mice undergo mutation and selection of high-affinity clones. **(a)** Pie charts indicating the frequency of the affinity-inducing mutation W33L in the CDR1 region of V_H_186.2 sequenced from single cell sorted NP+ IgG1+ CD38-GL7+ B220+ CD19+ cells from either young adult (7-10wo) or aged (90-100wo) B1-8i mice in recipient iLNs 14 days post-immunisation with NP-KLH/Alum. The values in the centre of the pie charts indicate the total number of cells sequenced per group (n=7-9 mice per group from two independent experiments). **(b)** Graph depicting the percentage of sorted GC B cells with the W33L mutation from young adult (7-10wo) or aged (90-100wo) B1-8i mice. Bar height corresponds to the mean, error bars indicate standard deviation, and each symbol represents values from individual recipient mice. **(c-d)** Graphs depicting **(c)** the number of single base pair mutations and **(d)** the ratio of replacement: silent mutations in the CDR1 region of V_H_186.2 among sorted GC B cells derived from young adult (7-10wo) or aged (90-100wo) at day 14 post-immunisation. Each symbol represents values from a single sorted GC B cell from 7-9 recipient mice per group. Statistics were calculated using unpaired Mann-Whitney U test. Data pooled from two independent repeat experiments.

## Discussion

In this study, we used an *in vivo* adoptive transfer system to show that B cells from aged mice have no intrinsic defects in affinity maturation post-immunisation, after they are transferred into young recipient mice. We first showed that B cells from aged mice had no intrinsic defects in class-switching to IgG1+ cells post-immunisation, consistent with previous work (Dailey et al., 2001; Lee et al., 2022; Russell Knode et al., 2019). Despite a lower proportion of follicular B cells among transferred NP-specific B cells from aged mice compared to those from young adult mice, NP-specific B cells from aged mice were not defective in becoming plasma cells or GC B cells, and had no difference in their DZ/LZ phenotype. The age-related reduction in percentage of follicular B cells was also reported in the spleens of aged mice (Turner & Mabbott, 2017). While we previously observed that B cells from aged SW_HEL_ mice preferentially entered the extrafollicular response and mounted a smaller early GC response compared to those from young adult mice (Lee et al., 2022), we did not observe this delayed kinetics in this system, suggesting some differences attributed to the model system used. Some factors that might contribute to this discrepancy include differences in antigen used that might result in distinct downstream BCR signalling following stimulation, and also differences in the organ analysed, spleen in the SW_HEL_ model and iLNs in the B1-8i model used here. Nevertheless, in both transfer systems, antigen-specific B cells from aged mice did not have intrinsic defects in mounting a peak GC response, implicating B cell extrinsic factors as causal in age-dependent defects in GC formation and magnitude.

Here we also showed that NP-specific GC B cells derived from B cells from aged mice were equally able to upregulate cMyc as those from young adult mice. cMyc expression is induced in GC B cells selected for their high-affinity BCRs which allows their cyclic reentry into the dark zone for further proliferation and somatic hypermutation (Calado et al., 2012; Dominguez-Sola, Victora, Ying, Phan, Saito, Nussenzweig, et al., 2012). Together with previous work showing no cell-intrinsic defects by B cells from aged mice in proliferating and upregulating costimulatory molecules after T-dependent stimulation *in vitro* and *in vivo* (Blaeser et al., 2008; Dailey et al., 2001; Lee et al., 2022), the lack of defects in cMyc upregulation shown in this study further supports the notion that B cells from aged mice are able to interact with and receive positive survival signals from Tfh cells and follicular dendritic cells normally. NP-specific GC B cells derived from B cells from aged mice also had similar rates of somatic hypermutation and frequencies of the affinity-enhancing W33L mutation as those from young adult mice. A previous study analysing the immunoglobulin genes of GC B cells from the Peyer’s patches and spleens of humans also revealed that the mechanism of somatic hypermutation is unaltered with age, although the same study reported variable changes in selection with age across organs, with an increase in selection in the GCs of spleen and decrease in Peyer’s patches (Banerjee et al., 2002). Further studies will be needed to fully dissect the effects of age on selection in different organs and immunisation regimens.

One limitation to our study is that the model system used, involving B cells with transgenic BCRs, is unable to account for B cell-intrinsic changes in the BCR repertoire, which has been shown to contract with age (Gibson et al., 2009; Tabibian-Keissar et al., 2016). Reconstitution of B10 scid mice with polyclonal B cells from aged mice was previously shown to result in lower mutation frequencies in the V_H_ sequences, compared to reconstitution with B cells from young mice (Yang et al., 1996). This might hint at a B cell-intrinsic defect in somatic hypermutation with age, although the number of sequences analyzed in the study (Yang et al., 1996) were significantly lower (17 to 40) than what was analyzed here. As such, whether age-related changes in BCR repertoire affect B cells’ ability to undergo affinity maturation remains to be characterised.

Our results collectively reveal that B cells from aged mice have no intrinsic defects in going through the cellular process that underpins affinity maturation in the GC. This implicates B cell-extrinsic factors as the key contributors to defects in the GC response and humoral immunity with age. Vaccine strategies aimed at improving vaccine responses in older people should therefore be targeted at improving the aged microenvironment, for example by rejuvenating Tfh differentiation and boosting stromal cell responses, to promote optimal B cell responses (Denton et al., 2022; Stebegg et al., 2020).

## Materials and Methods

### Mouse Husbandry and Maintenance

B1-8i BCR-Tg (Sonoda et al., 1997) and WT C57BL/6 mice were bred and maintained in the Babraham Institute Biological Support Unit (BSU), where B1-8i Tg mice were also aged. No primary pathogens or additional agents listed in the FELASA recommendations (Mähler et al., 2014) were detected during health monitoring surveys of the stock holding rooms. Ambient temperature was ∼19–21°C and relative humidity 52%. Lighting was provided on a 12 hr light: 12 hr dark cycle including 15 min ‘dawn’ and ‘dusk’ periods of subdued lighting. After weaning, mice were transferred to individually ventilated cages with 1–5 mice per cage. Mice were fed CRM (P) VP diet (Special Diet Services) ad libitum and received seeds (e.g. sunflower, millet) at the time of cage-cleaning as part of their environmental enrichment. All mouse experimentation was approved by the Babraham Institute Animal Welfare and Ethical Review Body. Animal husbandry and experimentation complied with existing European Union and United Kingdom Home Office legislation and local standards (PPL: P4D4AF812). Young adult B1-8i mice were 6-12 weeks old, and aged B1-8i mice were at least 90 weeks old when used for experiments. Young recipient C57BL/6 mice were 8-12 weeks old at the time of immunisation. All donor and recipient mice used in the experiments were female.

### Adoptive transfers of B1-8i cells

Single cell suspensions of spleen and mesenteric and peripheral lymph nodes from a young 6-12 week old adult and an >90 week old aged B1-8i mice were obtained by pressing the tissues through a 70 μm mesh in PBS with 2% foetal bovine serum under sterile conditions. B cells were then enriched using the MagniSort Mouse B cell Enrichment Kit (#8804-6827-74 Thermo Fisher Scientific), according to the manufacturer’s instruction. Cell numbers and viability were determined using a CASY TT Cell Counter (Roche). A small aliquot of enriched B cells was taken and stained to determine the percentage of NP-binding B cells by flow cytometry before cell transfer. The cell suspensions were then diluted in appropriate volumes of PBS to obtain a final concentration of 1 × 10^5^ NP-binding B cells/ml. 100μL of 1 × 10^4^ NP-binding B cells from young and aged donor B1-8i-Tg mice were injected intravenously into the tail of congenic WT recipients. Recipient mice were then immunised subcutaneously with NP-KLH/Alum, as detailed below, and draining inguinal LNs (iLNs) were collected at the indicated time points for flow cytometry.

### Subcutaneous immunisations with NP-KLH/Alum

To induce GCs in iLNs, recipient C57BL/6 mice were immunised subcutaneously on both flanks on the lower part of the body with NP-KLH (4-hydroxy-3-nitrophenylacetyl (NP)-Keyhole Limpet Hemocyanin (KLH), #N-5060-25 Biosearch Technologies). NP-KLH was first diluted in PBS and the same volume of Imject™ Alum (ThermoScientific #77161) was added to reach a final concentration of 0.5mg/ml NP-KLH. After 30 min of vortexing, 100μl of emulsion were injected subcutaneously (s.c.) into the hind flanks of the experimental mice, which have received the intravenous transfer of donor B cells.

### Phenotyping of B1-8i cells using Flow Cytometry

A small aliquot (1-2 × 10^6^) of enriched B cells from the young adult or aged donor mice was stained for phenotypic analysis prior to transfer. Briefly, cells were stained in 96-well v-bottom plates. Surface antibody staining was performed for 2 hours at 4 °C in PBS with 2% FCS, in the presence of 2.4G2 hybridoma (ATCC hb-197) tissue culture supernatant and Rat IgG isotype control (Invitrogen, #10700) to block non-specific binding via Fc interactions. Following incubation, samples were washed twice with PBS with 2% FCS, before they were fixed with the eBioscience Foxp3/Transcription Factor Staining Buffer (#00-5323-00) for 30 min at 4°C. The samples were then washed twice with 1 x Permeabilisation buffer (eBioscience #00-8333-56) and stained with the intracellular antibody mix in permeabilization buffer at 4°C overnight. Following overnight incubation, the samples were washed twice with 1 x Permeabilisation buffer and washed once with PBS with 2% FCS, before they are acquired on a Cytek Aurora Spectral Cytometer (Cytek). Cells for single colour controls were prepared in the same manner as the fully stained samples. The antibodies used for the surface and intracellular staining are listed in **Table 1**.

**Table 1.** List of antibodies used for flow cytometric analysis of B1-8i cells pre-transfer.

|  | **Antibodies used in Surface Stain** | **Company and Clone** | **Identifier** | **Dilution** |
| --- | --- | --- | --- | --- |
| 1 | BUV563-coupled anti-mouse CXCR4 | BD (2B11/CXCR4) | 741313 | 1:400 |
| 2 | BV480-coupled anti-mouse IgG1 | BD (A85-1) | 746811 | 1:500 |
| 3 | BV785-coupled anti-mouse CXCR5 | BioLegend (L138D7) | 145523 | 1:200 |
| 4 | PE-coupled NP | Biosearch Technologies |  | 1:100 |
| 5 | PE-Cy7-coupled anti-mouse CD40 | BioLegend (3/23) | 124622 | 1:2000 |
| 6 | A700-coupled anti-mouse MHCII IA/IE | Invitrogen (M5/114.15.2) | 56-5321-82 | 1:2000 |
| 7 | ViaKrome808-coupled Live/Dead | Beckman Coulter | C36628 | 1:2000 |
| 8 | APC-Fire750-coupled anti-mouse CD86 | BioLegend (GL-1) | 105045 | 1:1000 |
|  | **Antibodies used in Intracellular Stain** | **Company and Clone** | **Identifier** | **Dilution** |
| 9 | BUV395-coupled anti-mouse CD11c | BD (N418) | 744180 | 1:1000 |
| 10 | BUV496-coupled anti-mouse CD23 | BD (B3B4) | 741058 | 1:2000 |
| 11 | BUV661-coupled anti-mouse CD19 | BD (1D3) | 612971 | 1:2000 |
| 12 | BUV737-coupled anti-mouse CD62L | BD (MEL-14) | 612833 | 1:1000 |
| 13 | Pacific Blue-coupled IRF4 | BioLegend (IRF4.3E4) | 646418 | 1:1000 |
| 14 | BV510-coupled anti-mouse CD21 | BD (7G6) | 747764 | 1:2000 |
| 15 | AF488-coupled anti-mouse GL7 | Invitrogen (GL-7) | 53-5902-82 | 1:1000 |
| 16 | Spark YG593-coupled anti-mouse CD11b | BioLegend (M1/70) | 101282 | 1:2000 |
| 17 | Alexa Fluor 594-coupled anti-mouse Tbet | BioLegend (4B10) | 644833 | 1:500 |
| 18 | PE-Fire 640-coupled anti-mouse CD38 | BioLegend (90) | 102744 | 1:2000 |
| 19 | Spark NIR685-coupled anti-mouse IgD | BioLegend (11-26c.2a) | 405750 | 1:2000 |
| 20 | APCFire810-coupled anti-mouse B220 | BioLegend (RA3-6B2) | 103278 | 1:1000 |

### Flow Cytometric Analysis of iLNs post-immunisation

Cell suspensions of the dissected iLNs from the recipient mice were obtained by pressing the tissues through a 70 μm mesh in PBS with 2% FCS. Cell numbers and viability were determined using a CASY TT Cell Counter (Roche). Cells were washed and transferred into 96-well v-bottom plates. Surface antibody staining was performed for 2 hours at 4 °C in PBS with 2% FCS, in the presence of 2.4G2 hybridoma (ATCC hb-197) tissue culture supernatant and Rat IgG isotype control (Invitrogen, #10700) to block non-specific binding via Fc interactions. Following incubation, samples were washed twice with PBS with 2% FCS and acquired on a Cytek Aurora Spectral Cytometer (Cytek). Cells for single colour controls were prepared in the same manner as the fully stained samples. Flow cytometry data were analysed using FlowJo v10 software (Tree Star). The antibodies used are listed in **Table 2**.

**Table 2.**
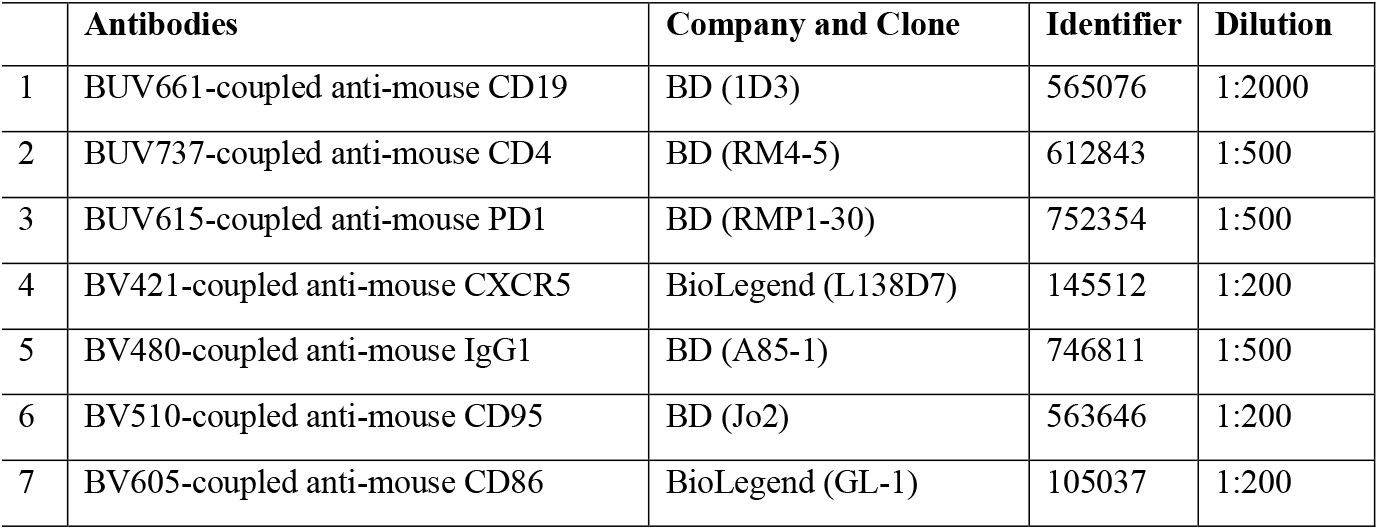
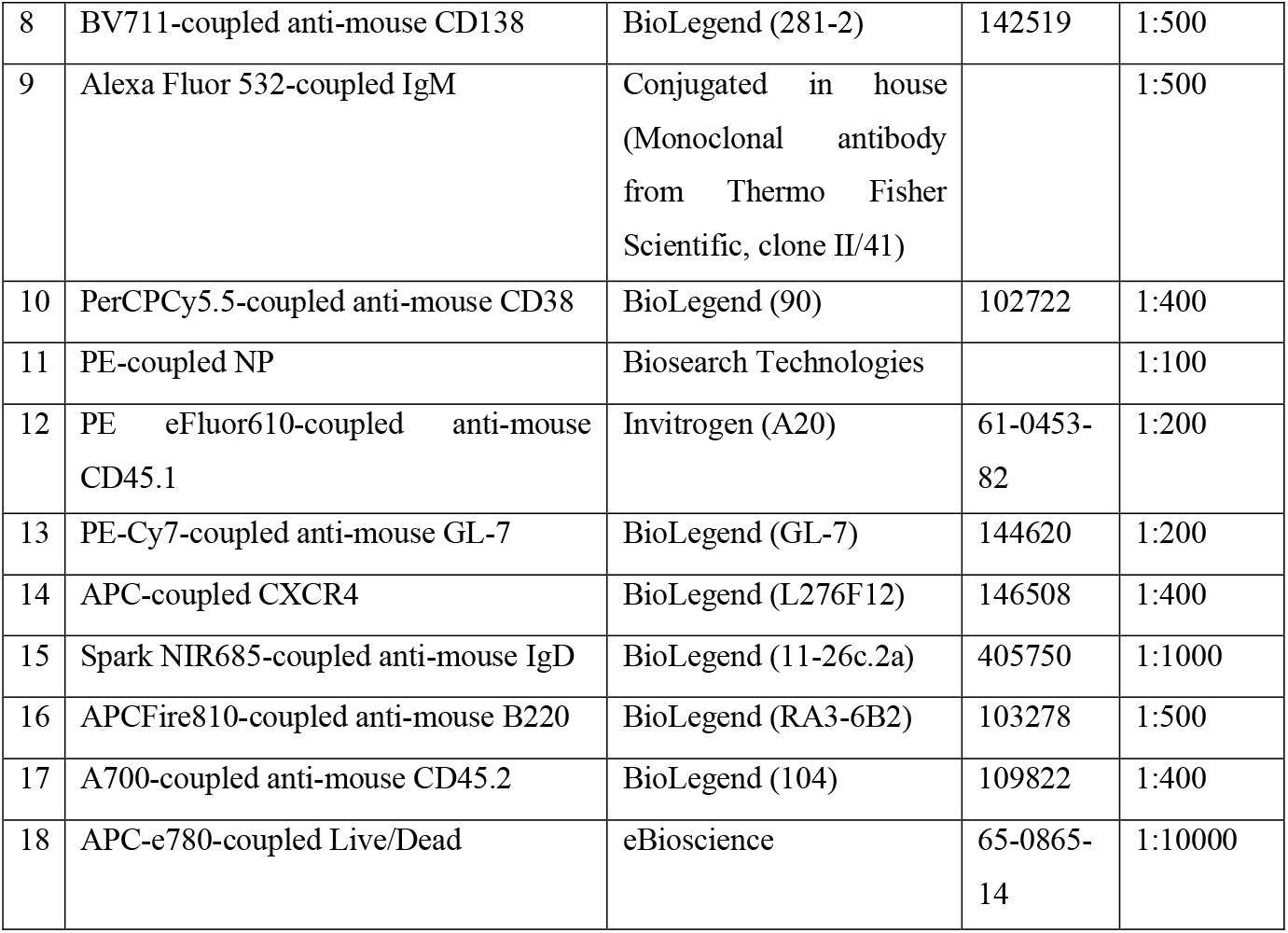
List of antibodies used for flow cytometric analysis of mouse LN cells post-immunisation.

### VH186.2 PCR and Sequencing of Single Cell Sorted GC B cells

Cell suspensions from iLNs were obtained as described above, and stained with the antibodies listed in **Table 3** for 2 hours at 4 °C in PBS with 2% FCS, in the presence of 2.4G2 hybridoma (ATCC hb-197) tissue culture supernatant and Rat IgG isotype control (Invitrogen, #10700) to block non-specific binding via Fc interactions. Anti-mouse CD3, CD4, CD11c, Ly6c antibodies were included in the DUMP APC channel to gate out non-B cells. After incubation, samples were washed twice with PBS with 2% FCS, filtered and resuspended.

**Table 3.** List of antibodies used for fluorescence-activated cell sorting (FACS) of mouse cells.

|  | Antibodies | Company and Clone | Identifier | Dilution |
| --- | --- | --- | --- | --- |
| 1 | BUV661-coupled anti-mouse CD19 | BD (1D3) | 565076 | 1:500 |
| 2 | BV510-coupled anti-mouse CD45.2 | BioLegend (104) | 109838 | 1:500 |
| 3 | BV605-coupled anti-mouse IgG1 | BD (A85-1) | 563285 | 1:500 |
| 4 | BV785-coupled anti-mouse B220 | BioLegend (RA3-6B2) | 103246 | 1:500 |
| 5 | PerCPCy5.5-coupled anti-mouse CD38 | BioLegend (90) | 102722 | 1:500 |
| 6 | PE-coupled NP | Biosearch Technologies |  | 1:100 |
| 7 | PE-Cy7-coupled anti-mouse GL-7 | BioLegend (GL-7) | 144620 | 1:500 |
| 8 | APC-coupled anti-mouse CD3 | BioLegend (17A2) | 100236 | 1:500 |
| 9 | APC-coupled anti-mouse CD4 | eBioscience (GK1.5) | 17-0041-83 | 1:500 |
| 10 | APC-coupled anti-mouse CD11c | BioLegend (N418) | 117310 | 1:500 |
| 11 | APC-coupled anti-mouse Ly6c | Invitrogen (HK1.4) | 17-5932-80 | 1:500 |
| 12 | A700-coupled anti-mouse CD45.1 | BioLegend (A20) | 110724 | 1:500 |
| 13 | APC-e780-coupled Live/Dead | eBioscience | 65-0865-14 | 1:10000 |

CD45.1+ CD45.2-NP+ IgG1+ B220+ CD19+ CD38-GL7+ cells were single cell sorted into 96-well plates, containing 10μl of reverse transcription lysis buffer per well (2U/μl RNase inhibitor (#EO0381 Thermo Fisher Scientific), 4mM DTT (#43816 Sigma), 30ng/μl Random Hexamers (#SO142 Thermo Fisher Scientific), 1% NP40, 0.2xPBS), using the FACSAria™ Fusion sorter (BD Biosciences). Plates containing the sorted cells were then stored at -80^0^C. For reverse transcription, plates were placed in a thermocycler (BioRAD), where it is heated to 65 °C for 2min and cooled to 10 °C for 5min. 15μl of reverse transcription buffer containing 1mM dNTPs (#R0194 Thermo Fisher Scientific), 8mM DTT (#43816 Sigma), 0.2U/μl RNase inhibitor (#EO0381 Thermo Fisher Scientific) and 160U GoScript Reverse Transcriptase (Promega #A5004) was then added to each well. The thermocycling program is then continued, with the following settings: 22° C for 10min, 37° C for 30min, 90° C for 6min. The prepared cDNA was stored at -20 ° C.

For nested PCR, 2.5μl of the prepared cDNA was used. The PCR mix for the first round of PCR was made up using 10x PCR buffer, dNTP and Taq DNA polymerase from the HotStar Taq DNA polymerase kit QIAGEN(#203205) with 20 pmol of the following primers: forward-GCTGTATCATGCTCTTCTTG and reverse-GGATGACTCATCCCAGGGTCACCATGGAGT. Plates were placed in a thermocycler (BioRAD) with the following program: 95° C for 15min, 94 °C for 3min, 39 cycles of 94° C for 45sec, 50° C for 1min and 72° C for 1min, and finally 72° C for 10min. The PCR product was then diluted 30 times in nuclease-free water. 1μl of the first PCR product was then used in the second round of PCR which was prepared with the HotStar Taq DNA polymerase kit (#203205 QIAGEN) and 20pmol of the following primers: forward-GGTGTCCACTCCCAGGTCCA and reverse-CCAGGGGCCAGTGGA T AGAC. The thermocycling program used for the second PCR was the same as the first. 5μl of the second PCR product was used to verify positive clones on a 1% agarose gel. The PCR products containing positive clones were purified using the ExoSAP-IT ™ PCR Product Cleanup Reagent (#78201 Applied Biosystems) and purified samples were sent for Sanger sequencing to Source Bioscience, UK. Analysis was performed using an automated alignment pipeline that aligned sequences to the VH186.2 sequence and the frequency of the affinity-enhancing W33L mutation was identified for each sequence, using an automated sequence trace analysis in R. Sequence traces were read using the *readsangerseq* function with default parameters from the *sangerseqR* package. Sequences shorter than 300 nucleotides or containing more than one N base call were omitted. The W33L locus was identified using *matchPattern* function from the Biostrings package, where it searches for ACCAGCTACTNNATGCACTGG in the reverse complemented sequence data while allowing 2 nucleotide mismatches and indels. A W or L call was assigned based on the following: if TNN in the appropriate position in the nucleotide string above was TGG the assignment was W, if TNN was TTA or TTG, this was assigned L, any alternative sequences for TNN were assigned other. Per sample calls were exported as a .csv for downstream analysis in the Prism v9 software (GraphPad).

## Statistical Analysis

All experiments were performed twice or more (3-5 recipient mice per group). Statistical analysis was performed using the Prism v9 software (GraphPad). Differences between experimental groups were determined using paired Wilcoxon matched pairs signed rank test, unpaired Mann-Whitney *U* test or 2-way ANOVA with Sidak’s multiple comparisons test, where appropriate. p values were considered significant when <0.05.

## Acknowledgements

We acknowledge the contribution of the Babraham Institute Biological Support Unit staff, who performed the *in vivo* treatments of our animals and took care of animal husbandry. We thank the staff of the Babraham Flow Cytometry for their technical support. This study was supported by funding from the Biotechnology and Biological Sciences Research Council (BB/W001578/1, BBS/E/B/000C0407, BBS/E/B/000C0427 and the Campus Capability Core Grant to the Babraham Institute). J.L.L is supported by a National Science Scholarship (PhD) by the Agency for Science, Technology and Research, Singapore. S.M.G is supported by the European Union’s Horizon 2020 research and innovation program “ENLIGHT-TEN+” under the Marie Sklodowska-Curie grant agreement No.: 955321. M.A.L. is an EMBO Young Investigator and a Lister Institute Prize Fellow.

## Author Contributions

J.L.L. designed the study, performed experiments, analyzed data, and wrote the manuscript. S.I., A.S.C., S.M.G. assisted with experiments. M.A.L. designed the study, wrote the manuscript, obtained funding, and oversaw the project. All authors read and edited the manuscript and contributed intellectually to this study.

